# A massive community-science dataset reveals convergent evolution of delayed flowering phenology in North American red-flowering plants

**DOI:** 10.1101/2024.09.25.614826

**Authors:** Patrick F. McKenzie, Andrea E. Berardi, Robin Hopkins

## Abstract

The radiation of angiosperms is marked by a phenomenal diversity of floral size, shape, color, scent, and reward. Through hundreds of years of documentation and quantification, scientists have sought to make sense of this variation by defining pollination syndromes. These syndromes are the convergent evolution of common suits of floral traits across distantly related species that have evolved by selection to optimize pollination strategies. The availability of community-science datasets provides an opportunity to develop new tools and to examine new traits that may help further characterize broad patterns of flowering plant diversity. Here we test the hypothesis that flowering phenology can also be a pollination syndrome trait. We generate a novel flower color dataset by using GPT-4 with Vision (GPT-4V) to assign flower color to 11,729 North American species. We map these colors to 1,674,908 community-scientist observations of flowering plants to investigate patterns of phenology. We demonstrate constrained flowering time in the eastern United States for plants with red or orange flowers relative to plants with flowers of other colors. Red-and orange-colored flowers are often characteristic of the “hummingbird” pollination syndrome; importantly, the onset of red and orange flowers corresponds to the arrival of migratory hummingbirds. Our results suggest that the hummingbird pollination syndrome can include flowering phenology and reveal an opportunity to expand the suite of traits included in pollination syndromes. Our methods demonstrate an effective pipeline for leveraging enormous amounts of community science data by using artificial intelligence to extract information about patterns of trait variation.

## Introduction

Adaptation in response to selection for optimized pollination is fundamental to the incredible diversification of flowering plants.^1–4^ The multi-dimensional response to selection has generated the evolution of correlated suites of traits associated with particular pollination strategies or pollinator vectors across angiosperms. Although agents of selection are complex and the phenotypic responses are highly variable, these so-called pollination syndromes have been a useful organizational theme for understanding the convergent evolution of flower color, size, shape, smell, and reward.^5–9^ Here, we leverage community science data and create a computer-vision-powered data extraction pipeline to compile an enormous dataset of North American flowering plants. We use this dataset to explore pollination syndromes and test how an additional trait – flowering phenology – can extend our knowledge of what it means to be a “hummingbird flower.”

Pollination syndromes reflect convergent evolution of suites of traits across distantly related plants in response to selection to optimize pollination strategies. For instance, wind-pollinated (anemophilous) flowers typically lack visual or olfactory appeal, presenting simple, inconspicuous blooms without bright colors or scents.^9^ In contrast, species pollinated by bees or butterflies (entomophilous) feature conspicuous flowers with abundant pollen and nectar. These flowers often possess specialized landing areas highlighted by UV-absorbing pigments to attract their pollinators.^10^ Fly-pollinated (myophilous or sapromyophilous) plants may exhibit less vibrant flowers, ranging from those with accessible nectar and pollen to blooms mimicking the appearance and scent of decay or rotting meat, often in shades of maroon or brown with speckled patterns.^11,12^ Meanwhile, hummingbird-pollinated (ornithophilous / trochilophilous) flowers are typically bright red or orange, tubular in shape, wide, have exserted stamens, and produce lots of nectar, catering to the feeding preferences of their avian pollinators.^2,5,7^ These pollination syndromes are not universal and represent simplifications of broad patterns; nevertheless, they can reveal dominant selective forces shaping reproductive strategies in flowering plants.^8^

As with any attempt at finding general patterns in the complexity of biological life, the ability to test the utility of pollination syndromes and expand upon the generalities of syndromes is limited by data availability. Naturalists, evolutionary biologists, and botanists have been compiling data over hundreds of years to formulate hypotheses about these syndromes, and yet our datasets are small relative to the enormous diversity of natural variation. Efforts to organize and standardize data across experiments have resulted in some useful datasets such as the TRY Plant Trait Database.^13,14^ Although this database is very large and has been actively curated for over 15 years, it is still limited in its scope – for example it only contains flower color data for around 2,500 of the 15,000-20,000 flowering plant species in North America. Recently, datasets that are orders of magnitude larger have been collected through community science projects. Community science has provided the general public with platforms, such as iNaturalist (www.inaturalist.org), to document natural diversity. Yet these datasets often lack curation, formatting, and trait specifications necessary for broader scientific inquiry. For example, out of over 44 million iNaturalist observations of flowering plants (as of September 2024) in our North

American study area, only roughly half (53.7%) have “research-grade” (a label assigned by iNaturalist for observations that have been reviewed and agreed on by multiple identifiers) species identifications. Despite these limitations, the rise of community science platforms offers a promising avenue for accessing large volumes of biodiversity data. This wealth of unstructured data presents both a challenge and an opportunity for expanding our understanding of pollination syndromes.^15^

The charismatic traits specifically associated with pollination syndromes, such as color, are ideal types of traits to mine from community science datasets. Color is among the most noticeable phenotypes to human observers, and perhaps unsurprisingly, some pollination syndromes are associated with a specific color or set of colors. For example, yellow flowers tend to attract generalist pollinators, blue flowers often attract bees, and flowers pollinated at night (e.g. by bats or hawkmoths) tend to be white.^9,16^ These associations are thought to often be driven by visual differences among pollinators: for example, many insects cannot see red and therefore have innate preference for other colors.^5,9,17,18^ In contrast, hummingbirds can see in the long wavelength spectrum and tend to specialize on red and orange flowers.^5,9,17–19^

Unlike flower color, one trait not typically included in pollination syndromes is flowering time.^20^ However, limited studies have suggested that pollinator availability may broadly shape where and when flowers of different colors occur. There are biogeographical patterns in some flower colors: for example, white and yellow flowers are the predominant phenotype in the arctic, where most pollinators are bees and flies.^9,21,22^ There are time-of-day patterns for some cases: for example, white-flowered species tend to open at night when crepuscular (i.e., active during twilight) or nocturnal pollinators such as hawkmoths and bats are active, and a previous case study ties flowering phenology to hummingbird presence for an Andean plant species.^9,23^ However, it is still unknown if seasonal flowering phenology is more widely correlated with pollination syndromes and related traits such as flower color.

We hypothesize that phenology is a key component of pollination syndromes, particularly for red-and orange-flowering plants associated with hummingbird pollination. North America is the ideal geographic region in which to test this hypothesis because of the spatial and temporal structure of hummingbird movement in this region. The ruby-throated hummingbird, *Archilochus colubris*, is the only species of hummingbird native to the eastern United States (with the exception of rare vagrants). They spend their winters in Central America and then migrate northward from March through May, where they breed, before migrating back to Central America in the late summer.^24^ In contrast, hummingbirds are largely absent from the middle of the United States, and several species (e.g. black-chinned, Anna’s, and rufous hummingbirds) are present along the west coast of the United States, one of which (Anna’s) is present year-round.

We predict that if the hummingbird pollination syndrome includes flowering phenology, then the flowering times of red-and orange-flowering plants will be restricted in the eastern United States, where pollination is limited by hummingbird migratory patterns. We expect no such restriction in other regions of the U.S. or in species with other flower colors. Importantly for our predictions, we observe a lag in the arrival of hummingbirds in the eastern U.S. relative to the overall onset of spring flowering. Between week 5 and week 12 of the year, flowers bloom in the mid-to north-eastern U.S. without hummingbirds present. Given that red and orange flowers are associated with the hummingbird pollination syndrome, we expect that these colors will be depleted from this region during these early weeks of the year relative to other colors and that the blooming of these plants will instead coincide with the arrival of hummingbirds after the 12th week of the year.

To test our hypotheses, we developed a novel pipeline that overcomes the challenge of limited data availability by extracting large-scale trait data from community science observations using an advanced computer vision model. Specifically, we mine the community science platform iNaturalist alongside GPT-4 with Vision to efficiently analyze flower color patterns across the U.S. This approach enables us to generate the most comprehensive dataset on North American flower color to date. With this dataset, we test our hypotheses about hummingbird-associated flowering phenology and demonstrate a scalable method for analyzing large-scale phenotypic patterns in community science data.

## Results

### Generation of a North American flower color database

We created a database of flower color for 11,729 North American flowering plant species by combining observation data from the community-science website iNaturalist and GPT-4 with Vision (GPT-4V) (pipeline summarized in Figure S1). This database includes 1,674,908 color-labeled observations, each of which is annotated as being in flower when observed and is associated with a time, location, and research-grade species identification. This final dataset was filtered from a total of 1,763,821 observations of angiosperms from iNaturalist, including 13,378 species, that were associated with the “flowering” phenology label as of October 11, 2023. GPT-4V was able to label the color of 88% of the species with only 1,637 species labeled as “NAN” or “UNKNOWN” flower color. With these species labels we were able to map flower color to 95% of the observations in our iNaturalist dataset. The cost of using GPT-4V for this task at this time was under 100 USD.

The size of our dataset is substantially larger than the TRY Plant Trait flower color dataset, which contains flower color for only 2527 species (2035 species overlapping with our dataset) in our study area. Our dataset is also a more accurate assessment of color as evaluated by human validation metrics. Specifically, out of a validation dataset of 250 species with a mix of flower colors, the three authors agreed with each other on 90% of the color labels (mean = 225, standard deviation = 3.60). The authors agreed with the GPT-4V color labels on 87% of species (mean = 217, standard deviation = 3) and agreed with the TRY color labels on only 81.6% of the species (mean = 204, standard deviation = 5.57). Furthermore, the authors visually inspected all 295 species labeled with red flowers from GPT-4V and were able to validate 85.4% of the species color assignments, representing over 95% of the red-labeled observations from iNaturalist (38,840 out of 40,742 observations). The mismatch between the percentage of species correctly labeled and the percentage of corresponding observations correctly labeled suggests that less-abundant species were more likely to be mislabeled.

The GPT-4V-labeled dataset had over 1 million more observations and over 10,000 more species than the TRY-labeled dataset, and more accurate color labels. Our query of iNaturalist also retrieved 163,693 observations for hummingbirds and 408,406 observations of bumblebees in North America for downstream analysis. These observations are similarly associated with time and location data.

### Patterns of flowering plants across North America

Our enormous GPT-4V-labeled dataset of flowering plants reveals general patterns in phenology across North America (Figure 1). Nearly 50% of both the species observed and total observations in the dataset belong to either the white or yellow color categories (Figure 1B). Red flowers only make up approximately 2.5% of the species and observations. Combining all colors, the average flowering time is consistent across all latitudes; however, the variance around the mean decreases dramatically with increasing latitude (Figure 1A). Lower latitudes tend to have a bimodal flowering pattern with a burst of blooming earlier in the year and a second burst late in the year, while flowering at higher latitudes is confined to a shorter season.

**Figure 1.**
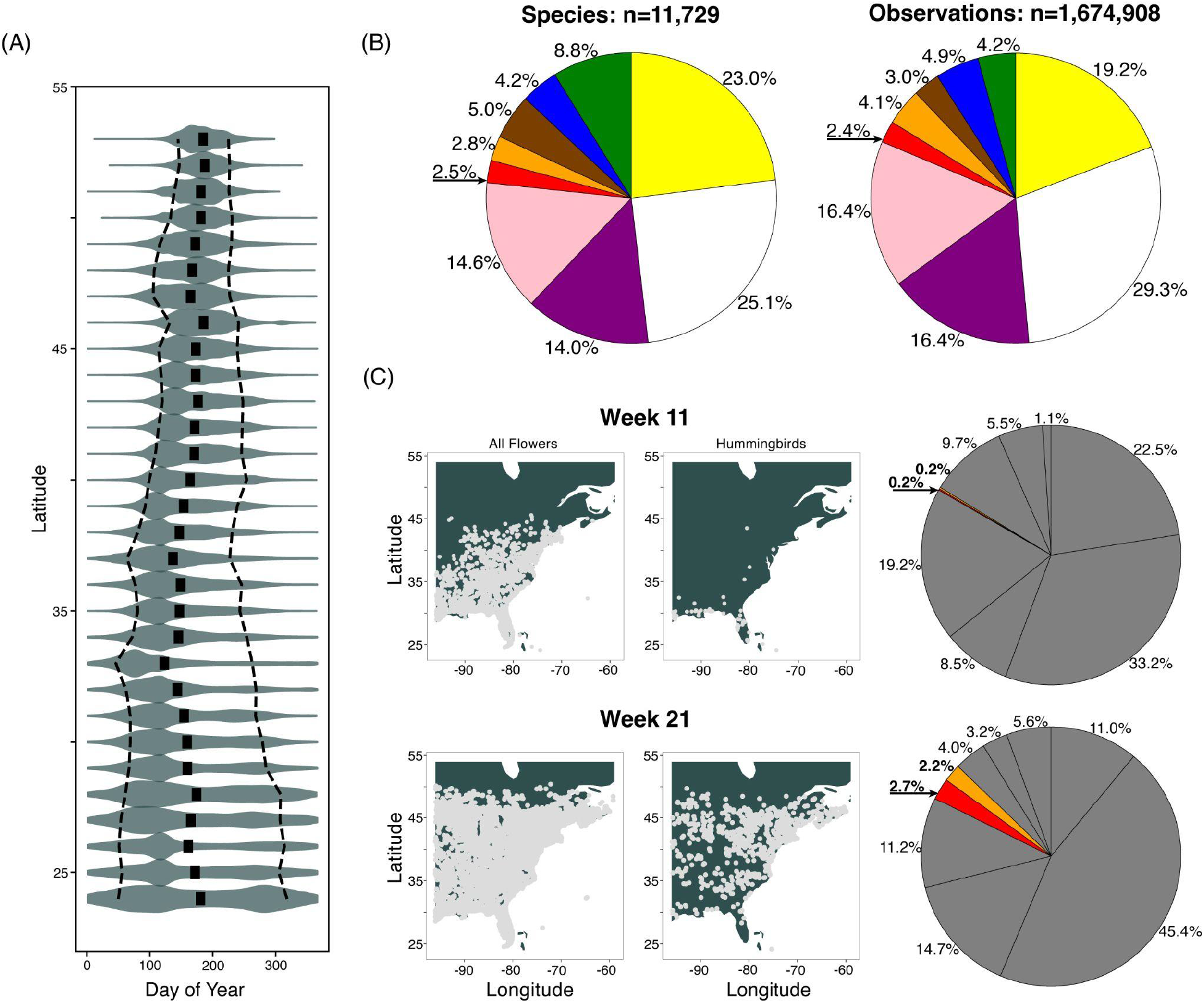
Summary of iNaturalist observation data. (A) Overall flowering phenology across all colors in the dataset with mean indicated by central black boxes, distribution indicated by gray violin plots, and 10th and 90th percentiles indicated by dashed black line. While the average flowering time is similar across latitudes, the variance of flowering times throughout the year decreases with increasing latitudes. (B) The flower color frequencies in the North American iNaturalist dataset show similar patterns when broken down by species (left) and number of observations (right). Colors clockwise from the black indicator arrow are: red, orange, brown/maroon, blue, green, yellow, white, purple, pink. The corresponding table can be found as Table S1. (C) Patterns of flowering plant and hummingbird observations in the eastern U.S. in week 11 of the year (top maps and chart) versus week 21 (bottom maps and chart). The maps show gray dots for every observation of flowering plants (left map) and hummingbirds (right map). In week 11, flowers are blooming throughout the eastern U.S., but hummingbirds have not yet arrived. By week 21, hummingbirds have migrated up the eastern U.S. The pie charts show a corresponding increase in the abundance of red and orange flower observations in the eastern U.S. by week 21 compared to week 11. Order of colors is the same as in Figure 1B. See Table S1 for full results.

As day of the year progresses from winter into spring, the number of flower observations increases, particularly in northern latitudes (Video S1). This general pattern of increasing observation data is also true for the two pollinators, bumblebees and hummingbirds, for which we also obtained data from iNaturalist. Observations of hummingbirds are strikingly absent from the eastern United States in the first weeks of the year and increase northward over time after the first three months of the year.

### Red-and orange-flowering plants bloom later than other colors

Flowering time of red- and orange-flowering plants lags behind other colors in the eastern half of the U.S. Red- and orange-flowering plants are conspicuously absent from the observation dataset in early spring, which corresponds to a period of time when hummingbirds are also not observed in this area. For example, in week 11 of the year, when hummingbirds have not yet migrated through the eastern U.S., red- and orange-flowering observations are scarce. In contrast, by week 21 when hummingbird observations are abundant, red and orange flowers are present and near their overall frequency in the dataset (Figure 1C).

The difference in flowering time can be visualized by tracking the northern edge (latitude of the northernmost 80^th^ percentile of flowering times) of flowering for plants of each flower color (Figure 2A). In the eastern U.S. (longitude -96 to -59), red- and orange-flowering plants start flowering 20-50 days after all other colors (white, blue, purple/pink, brown/maroon, green, and yellow). The lag begins about 20 days into the year, when flowers start blooming in the southeastern U.S., and the northern range of red- and orange-flowering plants catches up to the other colors around day 120.

**Figure 2.**
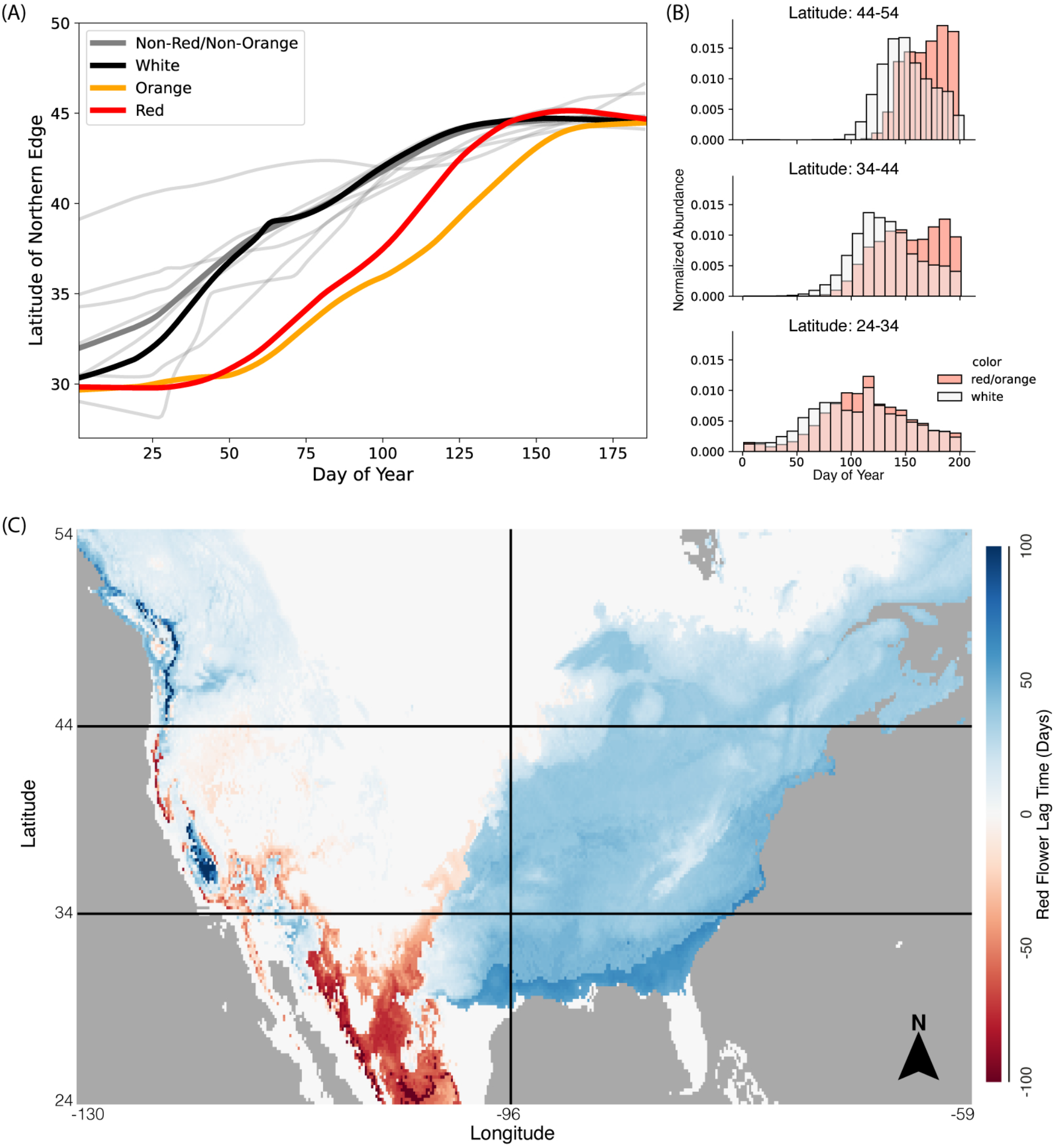
(A) Latitude of the northern edge (80th percentile of observations) of flower colors by day of year in the eastern U.S. Red and orange lag behind the other colors (shown in gray, with thicker gray line being the average of all other colors and thick black line representing white flowers). (B) Normalized abundances of red (in red) and white (in white) flowers for latitudinal sections of the eastern U.S. (24-34°N, 34-44°N, and 44-54°N) across the day of the year. (C) Consolidated results from MaxEnt maps corresponding to the difference in red and white flowering distributions in sliding windows through the year. The color of each pixel corresponds to the number of days by which white flower occupancy is inferred to occur before red flower occupancy for that pixel. Blue pixels indicate where white flowers occur before red flowers.

The pattern of delayed flowering in red and orange flowers is robust to overall abundance of each flower color observed. There are more white flower observations than red flower observations; to ensure patterns of flower time observations are not confounded with patterns of overall abundance we also compare red and white flowering abundance normalized to total abundance for each region of the study area (Figure 2B). As expected, the flowering season in the northern latitudes begins later than in the southern latitudes, as reflected in rightward movement of the histograms for the northern latitudes. In the eastern U.S., white flowers start blooming earlier than red flowers (the observation histograms for white flowers extend farther to the lower values on the left compared to red flowers), particularly at middle and upper latitudes (34N to 54N).

We quantified the extent of the flowering delay for red-flowered species across North American by modeling flowering time of red- and white-flowering observations only. We used MaxEnt distribution modeling software to infer the probability of occurrence of flowering by either white or red flowered plants for every pixel across North America for every day of the year. We defined the first day of flowering as the first day of the year where the model inferred the probability of occupancy for a flower color over a 0.5 threshold. The difference between red and white flowering times is extensive (heatmap shading in Figure 2C; Video S2). On average, white flowers are present 25.5 days (median=31.0, std=18.0) before red flowers broadly across the eastern U.S. In Mexico, red flowers appear to lead white flowers; however, there was very limited data associated with the high-elevation regions where this pattern was prominent, and white flower presence in some areas never passed the probability threshold that we used to determine occupancy (these cells take the darkest color in Figure 2C).

### Pollinators associated with flower color phenology differences

Red flowering time tracked the observations of hummingbirds but not bumblebees. Hummingbirds are known red-flower specialist pollinators, while bumblebees are considered to be broadly generalist pollinators. We found a striking overlap between the distributions of red flowers and hummingbirds (Figure 3A), as well as between white flowers and bumblebees. The latitude range for white flowers and for bumblebees advances northward earlier in the spring (beginning around days 20-40) than for red flowers and hummingbirds, both of which advance steeply later in the spring (beginning around days 60-80).

**Figure 3.**
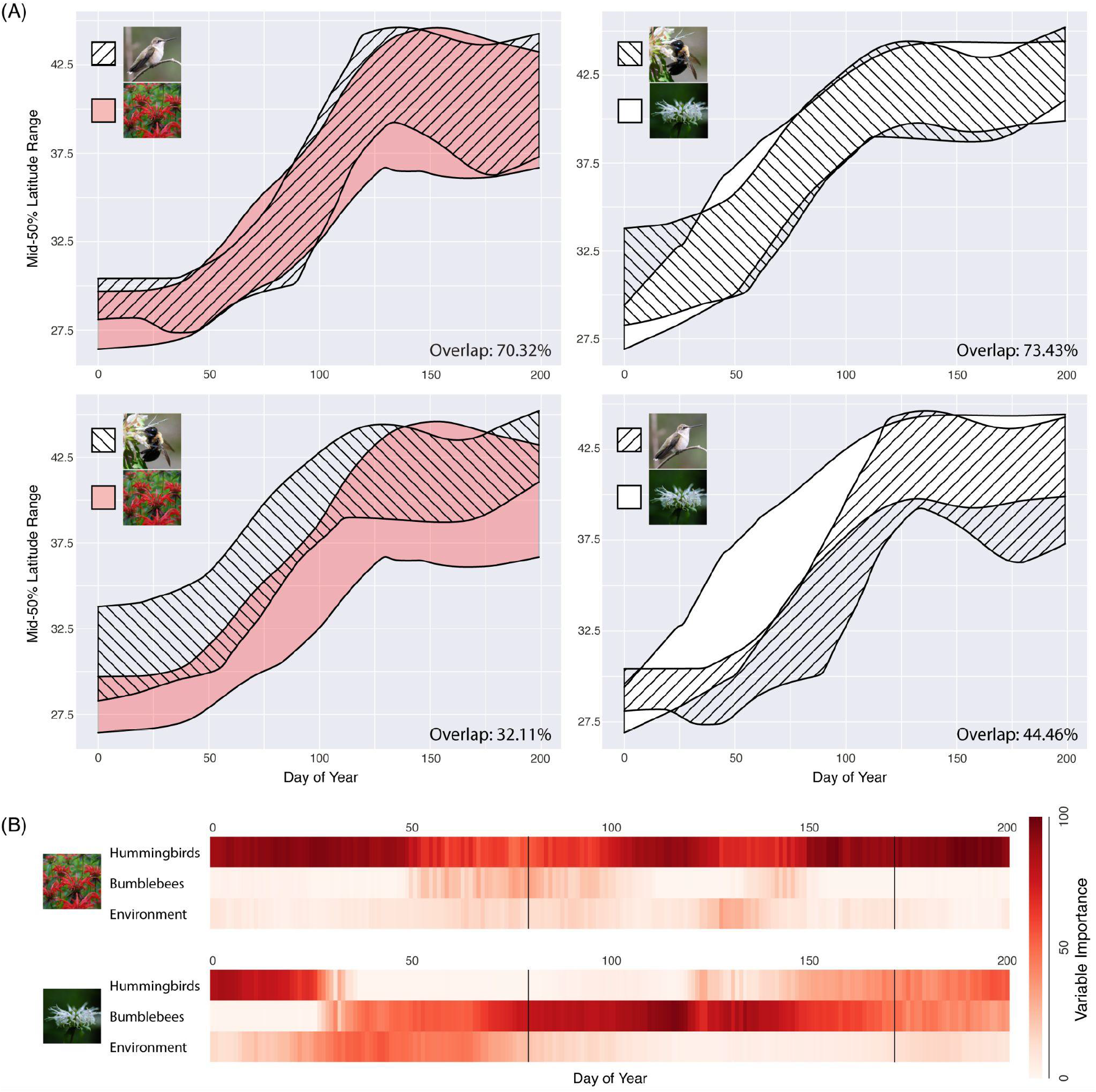
(A) The middle 50% latitude occupancy of red flowers, white flowers, hummingbirds, and bumblebees by day of year within the study area. On the top row, red flowers and hummingbirds are layered and white flowers and bumblebees are layered. The bottom row highlights pollinator mismatch, with bumblebees layered on red flowers and hummingbirds layered on white flowers. The average percent overlap across all days is reported in the bottom-right corner of each plot. (B) Importance assigned to each variable in MaxEnt distribution models for inferring the niche of red flowers and white flowers in 15-day overlapping windows through the year. Variables used for niche inference were seven environmental variables, as well as the inferred hummingbird distribution for that window and the inferred bumblebee distribution from that window. Black dividing lines indicate divisions between seasons. Darker red indicates stronger inferred importance for predicting flowering distribution.

We confirmed these observations by quantifying the degree to which pollinators can predict the presence of each flower color. We ran species distribution models in MaxEnt to find the distribution of each pollinator, hummingbirds vs. bumblebees, for each day of the year (in 15-day sliding windows). We then used these modeled distributions, along with environmental variables, to fit species distribution models for red flowers and white flowers. We found that for most of the year, hummingbird distribution is the most important predictor variable for red flower occurrence (Figure 3B top panel). In contrast, for most of the year, and most importantly during the expansion northward in the springtime, bumblebee distribution is the most important predictor variable for white flower occurrence (Figure 3B bottom panel).

## Discussion

We have created a novel pipeline using GPT-4V, a flexible artificial intelligence tool, alongside community science data to compare the flowering phenologies of different colors of flowering plants. We demonstrate that red- and orange-flowering plants in eastern North America bloom later than those of other colors and coincide closely with the seasonal arrival of hummingbirds, which preferentially visit red and orange flowers. Our results suggest that relative phenology can, in some circumstances, be included as a pollination syndrome trait, expanding our understanding of plant-pollinator interactions.

We compiled a large dataset of over 1 million iNaturalist observations from more than 10,000 North American flowering plant species. Each observation included a species identification, geographic coordinates, flowering status, and assigned flower color. Our use of GPT-4V for categorizing flower colors was highly successful, achieving accuracy comparable to inter-author consistency and outperforming a well-known plant trait database.

While our approach offers significant advantages, we acknowledge the limitations inherent in using community science data. Longstanding methods for precisely measuring floral color – e.g., pigment extraction, spectral reflectance analysis, and standardized photography – are essential for achieving fine-grained accuracy but limit the scale at which floral color data can be analyzed.^25,26^ By using community science data together with computer vision, we traded some precision for quantity, enabling us to construct a dataset quickly and at minimal cost, while still maintaining high accuracy at the level of coarse color categories. To our knowledge, this is the most comprehensive flower color dataset ever assembled for a geographic region of this size. We also recognize known biases associated with iNaturalist data, such as observations being concentrated in urban areas and showier flowers tending to be disproportionately represented.^27^ However, since we are primarily examining broad, latitudinal flowering patterns within each color group, we do not expect these biases to significantly impact our results. Future research might incorporate herbarium specimens, which exhibit less taxonomic and geographic bias than iNaturalist data.^28^

Previous studies have explored machine learning approaches for extracting color and other information from plant image data, including from community science datasets.^29–33^ However, these methods have been limited by the need for extensive model training and manual identification of the flowers within images, which restricts their scalability. In contrast, large multimodal models (LMMs) like GPT-4V offer the advantage of generality due to their pre-training on diverse datasets, eliminating the need for specialized training and manual annotation. This generality enabled us to perform extensive sampling, enhancing the statistical power of our analysis. Nevertheless, the LMM approach may not be universally applicable. Targeted studies focusing on flower color in a more constrained set of taxa or using photos with consistent composition and exposure might achieve finer-grained characterization. In such cases, dedicated models specifically trained for these purposes could better capture subtle differences in floral color, with biases that are easier to characterize and account for.

By capitalizing on our novel methodology, our study reveals biological patterns that deepen our understanding of pollination syndromes. We focused on comparing red- and white-flowering plants because their colors are easily distinguished, and they typically attract different pollinators. By demonstrating that flowering time in red-flowering plants lags behind that of white-flowering plants in the eastern U.S., our analysis underscores the influence of pollinator availability on flowering phenology. While the phenology of white flowers corresponded well with that of other colors in our analysis of North American plants, other patterns in the data remain to be explored – for instance, the apparent early appearance of maroon/brown flowers in the eastern U.S. relative to other colors (Figure 2A, top-left gray line).

These findings suggest that the convergent evolution toward hummingbird pollination extends beyond the traditional physical floral traits (red, tubular flowers with exserted stamens) associated with the hummingbird pollination syndrome. Specifically, we observe that in the eastern U.S., flowers expected to be hummingbird-pollinated bloom later in the spring than those of any other color, indicating that flowering phenology itself might be a facet of the hummingbird pollination mode, particularly in locations where hummingbird presence is seasonally constrained. With climate change causing hummingbirds to arrive earlier at northern latitudes, flowering phenology may come under strong directional selection to maintain synchrony with pollinator availability.^24,34^ Although our analysis focuses on a single example of phenology-related constraints, future analyses could extend this approach by examining other seasonally constrained pollinators, exploring different geographic regions, or investigating additional physical traits associated pollination syndromes (e.g., floral shape instead of floral color). Such research is essential for refining our understanding of the conditions under which flowering phenology functions as a pollination syndrome trait.

Future work at the intersection of phenology and community science data could further scale up our data extraction approach to include more observations and/or additional traits. In creating our dataset, we used a representative photo from each species to assign color to the entire species, which excludes within-species variation. Reducing the task from labeling each *observation* to labeling each *species* saved money, keeping the API use cost under 100 USD, and was also necessitated by the rate limits of the GPT-4V preview model. As computer vision models become more affordable and allow higher-throughput analysis, future approaches might instead query each individual observation. This would account for possible color polymorphism and enable us to assign flower color to a species based on a summary across many observations. It would also eliminate the dependence on manually annotated “flowering” phenology through iNaturalist, as recent work has shown that computer vision could directly determine whether the plant in each observation is flowering.^35^

We anticipate that our iNaturalist-to-GPT-4V pipeline can be generalized to study other aspects of floral phenotypes observable in community science data, such as investigating replicated patterns of within-species floral variation. Additionally, we envision studying color in a more fine-grained manner by treating color assignments as continuous variables in, for example, RGB space. While extracting these values meaningfully from community science photographs is challenging due to the lack of standardized color calibration across images, advancements in automated photo calibration tools and computer vision models may soon overcome these hurdles. Harnessing these emerging technologies alongside massive community science datasets like iNaturalist could unlock unprecedented insights into plant phenotypic diversity, transforming our understanding of global patterns of plant variation.

## Materials and Methods

### Study area

We limited our study area to the contiguous lower United States, northern Mexico, and southern Canada with a latitude/longitude-bounded box ranging from -130 to -59 degrees in longitude, and from 24 to 54 degrees in latitude.

### iNaturalist observation data

We used the iNaturalist data export tool (https://www.inaturalist.org/observations/export) to collect data from our study area for all flowering plants (Angiospermae). We filtered the export to only include observations that were labeled as being in flower, had open geoprivacy, had photos, and whose identifications were marked as research grade. For analysis of hummingbird and bumblebee distributions, we also exported all observations of each of those groups (family Trochilidae and genus Bombus, respectively) that had photos, had open geoprivacy, and were research grade.

### GPT-4V flower color data

From the outset we intended to produce a large dataset of observations of flowers matched to their corresponding flower colors. To maximize the size of our dataset, we employed a new approach for bulk labeling of flower color, using a computer vision model from generative AI. Our approach was to assign a flower color for each of the species in the dataset and then to project these species-assigned colors back onto the full dataset of observation data (see Figure S1). To do this, we: 1) used the iNaturalist API to match each species to a representative “default” photo from the iNaturalist website, and 2) used a cutting-edge general computer vision model to assign each image with a categorical label describing the color of the flower. We interacted with the iNaturalist API in Python through the client “pyinaturalist.” We queried each species individually using the “get_taxa” function and saved the returned “default_photo” url alongside each species into a two-column dataframe – this dataframe had species names in the first column and a link to each representative photo in the second column.

For the computer vision step, we used the OpenAI API, which allows us to programmatically interact with OpenAI’s suite of generative AI models in Python. We used the “GPT-4 with Vision” model, which in late 2023 – early 2024 was available in a rate-limited preview capacity, by designating the model choice as “gpt-4-vision-preview”. The GPT models are designed to be very general, and the vision model can interpret basic features of both text and photographs. For each query, we passed GPT-4V the url link to the representative species photo along with the following instructions:

“Please adhere to very specific formatting in your response: three words separated onto three lines (one word per line). The first line should indicate ‘YES’ or ‘NO’ to answer whether there is a flower present. The second line should be one word from the following list, to best describe the flower color in the photo: [‘BLUE’, ‘BROWN’, ‘GREEN’, ‘ORANGE’, ‘PINK’, ‘PURPLE’, ‘RED’, ‘MAROON’, ‘WHITE’, ‘YELLOW’,’UNKNOWN’,’NAN’]. The flowers might not match these categories perfectly. Do the best you can. If in doubt, please be conservative and choose ‘unknown’. The third line should indicate your assessment of the subjectivity of the answer --it should either be LOW, MEDIUM, or HIGH, where HIGH means that the choice of color assignment seems highly subjective.”

### TRY Plant Trait Database color data

To help benchmark our flower color inferences, we accessed all non-restricted flower color data through the TRY Plant Trait Database. To do this, we queried public data for the traits “flower color” (ID=207) and “corolla color” (ID=3866). Like our GPT-4V-labeled data, the TRY data was associated with species. Because it is a global dataset, we trimmed down the dataset to only those species present in the iNaturalist dataset from our study area.

### Validation

We conducted a validation analysis to ensure reliability in categorical color label assignments, focusing on the 250 species with the highest abundance that had flower color labels available in both the GPT-4V and TRY datasets. Each of the three authors manually scored flower color for all 250 species from the following pool of colors: blue, yellow, green, white, purple / pink / violet, orange, red, maroon / brown, and black. We averaged pairwise comparisons among the three authors with the GPT-4V and TRY labels. Given that multiple possible colors are associated with some species in the TRY data, we considered it a match if any of the TRY colors for a species corresponded with our assigned color. We also manually validated all species labeled as red.

### Analysis

We analyzed our data using a combination of R and Python notebooks. For the purposes of the analysis, we combined the color labels of pink/purple and brown/maroon.

### Comparing northern edges of each flower color

Our primary focus initially was to illustrate the relationship between time of flowering (converted to an integer “day of year” ranging from 0 to 365) and location, specifically latitude. We subset the data to eastern (-96W to -59W) North America. In this region, we recorded the latitude of the northern edge of the distribution of each flower color across the ordinal dates of the year, using a sliding window of 25 days. Specifically, for each color category and for each 25-day window, we extracted all observations from our dataset with their corresponding latitudes, and recorded the value 80th percentile using the R builtin quantile function. This gives us the northern latitude edge of flowering for each flower color for each 25-day period.

### Comparing red vs. white flower abundance

With the datasets of just red and white flower colors we summarize trends in flowering time by region for each color. We compared the relative abundance of observations of each color in different parts of North America. To do this, we subset the data to the eastern U.S. (-96W to -59W), and we further subset the data by latitude into 3 rows (24N to 34N, 34N to 44N, 44N to 54N). Within each grid cell we created histograms showing the normalized abundance of each color by ordinal day. This demonstrates the different flowering distributions of the two colors over space and through time.

### Quantifying lag in flowering time across the landscape

We used niche modeling in sliding windows to determine occupancy of flowering plants across the landscape for each flower color through time. Distribution modeling is often used to infer the broad distribution of where a species or group of species can or could occur given some observations of the species and patterns of environmental variables. Instead of inferring occupancy of a species, we inferred occupancy of a flower of a particular color. Separately for red and for white, we subset the “flowering” observations in overlapping 15-day windows, sliding one day for each window. For each data subset, we used MaxEnt species distribution modeling program (version 3.4.4) to incorporate the observation data along with seven WorldClim 2.0 (10-minute) environmental variables (annual mean temperature, maximum temperature of warmest month, minimum temperature of the coldest month, annual precipitation, precipitation of the wettest month, precipitation of the driest month, and elevation [from the Shuttle Radar Topography Mission, accessed through WorldClim]) to predict, cell by cell, the probability of occupancy of that particular flower color during that particular window. The study area was represented by a 180×426 matrix.

Once we estimated occupancy of flowering, we calculated the extent to which red and white flowers bloomed at different times. To do this, we identified the first day of flowering for each color for each pixel as the point at which the MaxEnt-output value of that pixel was greater than 0.5. With two matrices of “first day of flowering” values for red and white flowers separately, we then subtracted the white flower matrix from the red flower matrix to get a matrix for which each pixel represented the number of days by which white flowering time led that for red flowers. When graphed, this represents the difference in flowering phenology of red and white flowers across space. We calculated the mean, median, and standard deviation of cells of this difference matrix in the eastern U.S. to quantify the extent to which white flowers lead red flowers.

### Pollinator associations – hummingbirds and bumblebees

We tested the hypothesis that flowering time of red flowered plants is constrained in the eastern United States because this is where ruby-throated hummingbirds are seasonal migrants in the eastern United States. We investigated how well the presence of two different pollinators – hummingbirds, which are strongly associated with pollinating red flowers, and bumblebees, which are generalist pollinators that usually avoid red flowers – predicted the phenology of red and white flowers.^9,17,18,36^ For our study area, we exported all research-grade iNaturalist observations for bumblebees (genus *Bombus*) and hummingbirds (family Trochilidae). We used the pollinator data for two analyses: comparing the 25% (trailing edge) and 75% (leading edge) latitudes of red and white flower colors with the pollinators, and for niche modeling where the pollinator distributions were used as predictors.

For the first analysis, we subset the data to look at the eastern United States, where hummingbirds are widespread residents only during the late spring and summer. We used a sliding window each day of the year (starting edge of the window) and we calculated the 25% percentile latitude and the 75% percentile latitude. We repeated this across red flowers, white flowers, hummingbirds, and bumblebees, so that for any day of the year we have an estimate of latitude range. We then compared the distributions of flowering across latitude for these different flower colors and pollinators.

For the second analysis, we used MaxEnt for niche modeling. We first inferred distributions for hummingbirds and bumblebees separately for each day of the year, using 15-day sliding windows, linear models, and the seven predictor variables used for the flowering analysis above. Once we had these results, we then turned to red and white flowers and again ran MaxEnt for each day of the year. For these runs, we used 15-day sliding windows, linear models, and the previous seven “environmental” predictors, but we also included two more predictor variables for each run: the hummingbird and bumblebee distributions from the same window of the year. After these runs finished, we examined variable importance to determine whether environmental variables, hummingbird distributions, or bumblebees distributions were important in explaining red vs. white flower distributions for each day of the year.

## Supporting information

GPT-labeled species dataset

Validation of red-labeled species

## Acknowledgements

We are grateful to the community of iNaturalist users, curators, and staff for generating and maintaining the data underlying this project. Thanks to the Hopkins Lab and Deren Eaton for feedback on early versions of this project. R. H. was funded by NIH NIGMS-1R35GM142742-01 and NSF DEB-1844906.

