## Supplemental Figure 1, Supplemental Table 1 for "A massive community-science dataset reveals convergent evolution of delayed flowering phenology in North American red-flowering plants"

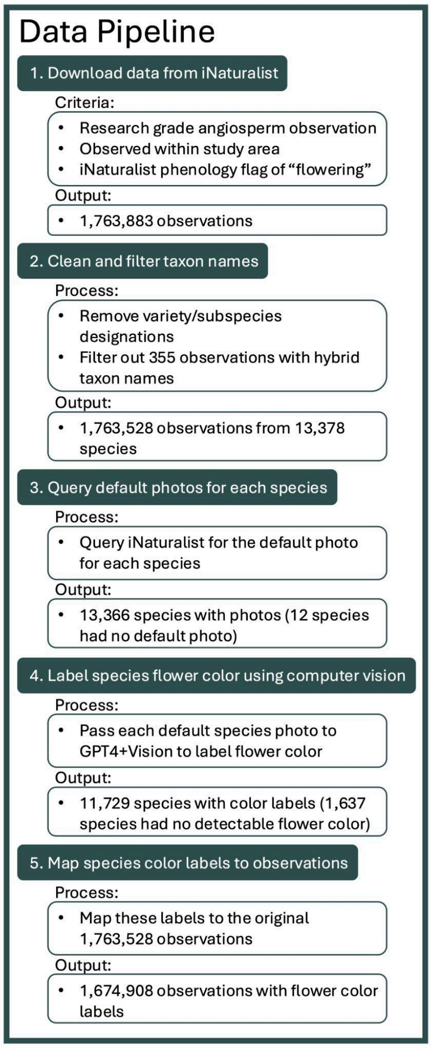


Figure S1: Data collection, cleaning, and annotation pipeline.

| color | percent_species | percent_occurrences | percent_occs_week11 | percent_occs_week21 |
|---|---|---|---|---|
| yellow | 23.0 | 19.2 | 22.5 | 11.0 |
| white | 25.1 | 29.3 | 33.2 | 45.4 |
| purple | 14.0 | 16.4 | 8.5 | 14.7 |
| pink | 14.6 | 16.4 | 19.2 | 11.2 |
| red | 2.5 | 2.4 | 0.2 | 2.7 |
| orange | 2.8 | 4.1 | 0.2 | 2.2 |
| brown/maroon | 5.0 | 3.0 | 9.7 | 4.0 |
| blue | 4.2 | 4.9 | 5.5 | 3.2 |
| green | 8.8 | 4.2 | 1.1 | 5.6 |

Table S1: iNaturalist occurrence data broken down by flower color. This table corresponds to the pie charts in Figure 1.
